# Increased maize chromosome number by engineered chromosome fission

**DOI:** 10.1101/2025.02.05.636704

**Authors:** Yibing Zeng, Mingyu Wang, Jonathan I. Gent, R. Kelly Dawe

## Abstract

Activation of synthetic centromeres on chromosome 4 in maize leads to its breakage and formation of trisomic fragments called neochromosomes. A limitation of neochromosomes is their low and unpredictable transmission rates due to trisomy. Here we report that selecting for dicentric recombinants through male crosses uncovers stabilized chromosome 4 fission events which split it into 4a-4b complementary chromosome pairs, where 4a carries a native centromere and 4b a synthetic one. The cells rapidly stabilized chromosome ends by de novo telomere formation and the new centromeres spread among genes without altering their expression. When both 4a and 4b chromosomes were present in a homozygous state, they segregated through meiosis indistinguishably from wild-type, and gave rise to healthy plants with normal seed set. This work leverages synthetic centromeres to engineer chromosome fission, effectively raising the diploid chromosome number of maize from 20 to 22.

**TEASER:** Using synthetic centromeres, we divide one maize chromosome into two chromosomes, and show that both chromosomes are fully functional.

## INTRODUCTION

Karyotype engineering is a form of genome editing that involves altering large segments of a genome, joining chromosomes, or potentially increasing the number of chromosomes ^1^. Creating a new chromosome is particularly challenging because it requires engineering a new centromere. Centromeres are composed of DNA and a large number of kinetochore proteins that interact with DNA to facilitate microtubule binding, sense proper chromosome alignment, and initiate anaphase ^2^. However, in most cases the interaction between centromeric DNA and kinetochore proteins is not sequence specific. Virtually any DNA sequence can serve as the structural foundation for a kinetochore ^3,4^, though sequences that are repetitive and gene-free are most common in nature ^5,6^. Once a centromere position is established, the replication process is epigenetic: specialized deposition proteins mediate the recruitment of additional kinetochore proteins to the same site over subsequent cell cycles ^7,8^.

The lack of sequence-specificity means that engineering a new centromere requires a method to recruit inner centromere proteins to a designated centromere platform. One method, called protein tethering, involves fusing centromere proteins to a DNA binding protein that recognizes sequence motifs in the designated centromere platform. Most or all of the known functions of the centromere can be recruited to a synthetic repeat structure by tethering the key centromere protein CENP-A/CENH3, chaperones that recruit CENH3, or several other proteins that interact closely with CENH3 ^9–14^. Synthetic centromeres formed through protein tethering have been shown to bind microtubules and confer mitotic chromosome segregation in multiple species ^10,14–20^. We recently demonstrated that synthetic centromeres can be induced in maize and transmitted through meiosis ^21^. We tethered a LexA-CENH3 fusion protein to an array of LexO binding sites on the long arm of maize chromosome 4, and showed that new centromeres cause the formation of dicentric chromosomes and chromosome breakage. Broken sections of chromosome 4 were recovered as trisomic neochromosomes (4b chromosomes) harboring new centromeres at the LexO array. The neochromosomes were propagated for several generations, but the transmission frequencies were low due to meiotic errors and gene dosage effects that accompany trisomic chromosomes.

Here, using synthetic centromere technology, we describe the fission and recovery of two chromosomes from a single ancestral chromosome. Starting with a trisomic 4b line, we demonstrate that crossing over between two centromere locations leads to dicentric derivatives that undergo breakage-fusion-bridge cycling followed by de novo telomere formation, and stabilization of a 4a-4b new karyotype with overlapping ends. We show that the expression of 22 genes embedded with the 4b centromere are not significantly different from the progenitor line, and that DNA methylation is unchanged in the intergenic spaces except for slight reductions in the CHG context immediately beneath CENH3. We further show that the synthetic centromeres support meiotic chromosome transmission that is indistinguishable from wild type.

## RESULTS

### Molecular structure of the ABS4 binding platform

Our system for engineering synthetic centromeres is based on a binding platform called Arrayed Binding Sites 4 (ABS4) that contains thousands of LexO binding sites on the long arm of chromosome 4. ABS4 was created by mixing PCR products containing a repeating 157 bp monomer with a marker plasmid (pAHC25) and transforming the mixture by biolistic transformation into the maize Hi-II line (which is related to the A188 inbred ^22,23^). For this study we self-crossed a line carrying ABS4 five times, and from one plant constructed a PacBio HiFi library and sequenced and assembled the genome.

The data show that the ABS4 inbred contains two insertions approximately ∼1.45Mb from each other. The first is only ∼7 kb, while the second is ∼583 kb. The larger insertion contains 467 kb of the ABS monomer interspersed with 117 kb of fragmented copies of the pAHC25 plasmid (Fig. 1), though we do not have a complete picture of the internal structure because there remains an assembly gap in this region. Within the assembled sequence there are approximately 2,800 copies of LexO. The transformation process also created a rearrangement of the native genome. A comparison of the ABS4 assembly to the A188 reference assembly ^24^ indicates that a portion (∼360 kb) of the genome next to the first insertion was inverted and inserted downstream of the second insertion.

**Figure 1.**
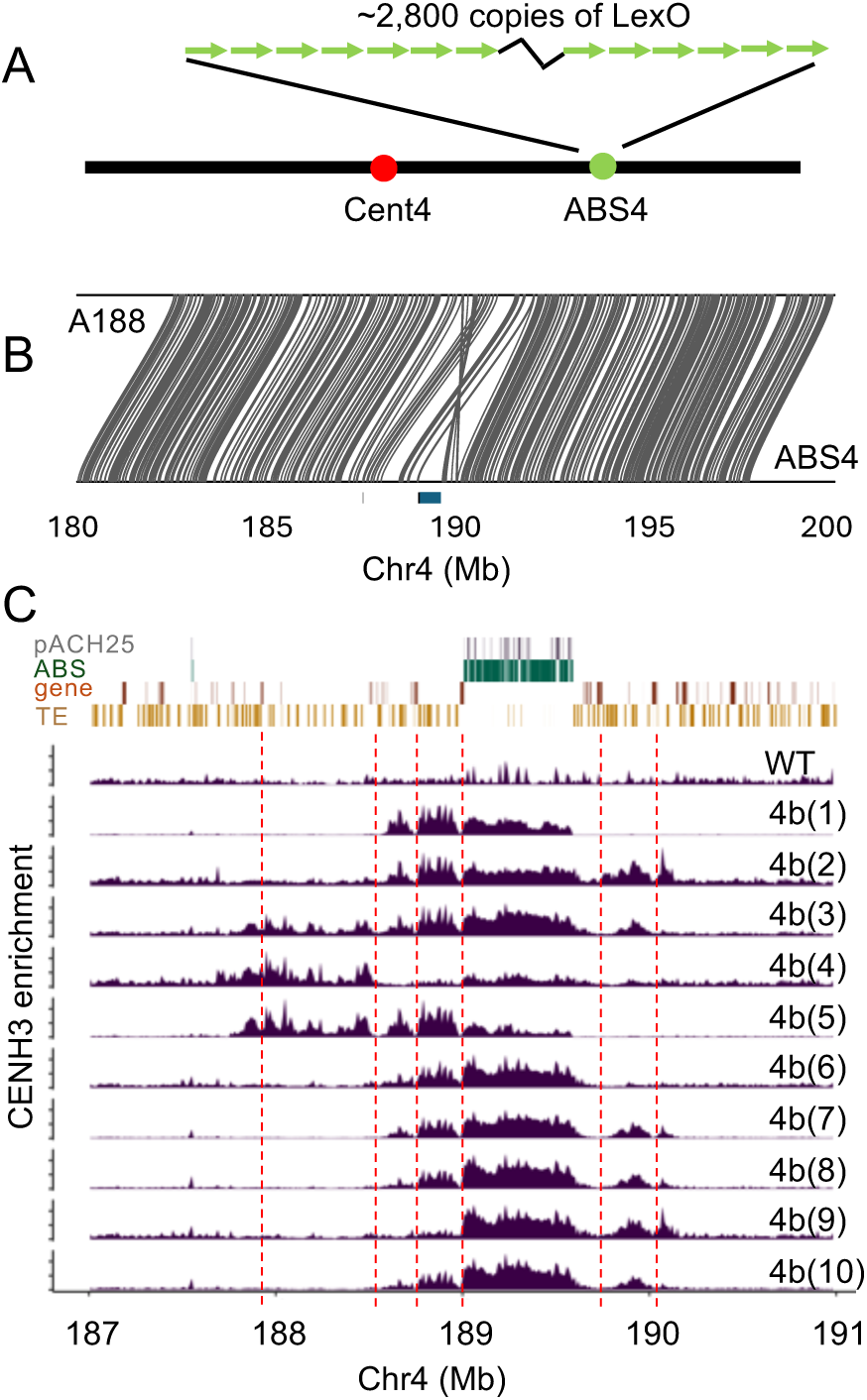
Newly formed centromeres over ABS4. **A**) Schematic of chromosome 4 showing the locations of normal centromere 4 (approximated by Cent4) and ABS4. **B**) Assembled region of the ABS4 genome and comparison with the progenitor chromosome from the inbred A188. **C**) Annotation features in the ABS4 region and CUT&Tag data from 10 neochromosomes. Red dotted lines show locations dips in CENH3 enrichment associated with genes.

### Selection for 4b neochromosomes

Our method for inducing centromeres at ABS4 involves crossing it to a line expressing a LexA-CENH3 fusion protein ^21^. When LexA-CENH3 binds to the LexO binding sites in ABS4, it recruits native CENH3 and sufficient overlying kinetochore proteins to form a second active centromere on chromosome 4. Dicentric chromosomes are unstable because the two centromeres frequently move to opposite poles in mitosis, creating a bridge during anaphase that is broken. The break can either be repaired by *de novo* telomere formation or by fusing with the broken end from the other chromatid In the next cell cycle. If the break is repaired by fusing with another broken chromatid, a dicentric chromosome is reformed, and the process can repeat itself in what Barbara McClintock referred to as the Breakage Fusion Bridge cycle (BFB)^25^. McClintock demonstrated that BFB can be maintained for multiple cell cycles before telomeres ultimately form (what she called “healing”) and halt the process ^26,27^. The mechanism of *de novo* telomere formation is not well understood, though at least in vitro, the maize telomerase enzyme can extend telomere repeats on a variety of DNA templates ^28^

When broken fragments formed during ABS/LexA-CENH3-activated BFB are healed by telomere formation, the outcome is a new chromosome (neochromosome) with an engineered centromere at ABS4. Our objective was to recover newly formed neochromosomes in progeny and study their behavior in subsequent generations ^21^. As described previously, we designed a screen to pass neochromosomes through the female gametophyte along with a complete chromosome 4. Instead of tracking ABS4 directly, we used alleles of a seed pigment gene called *colorless2* (*c2*) that is linked to ABS4 by about 15 cM. The female parent was heterozygous for ABS4 and the dominant *C2* allele that pigments seeds (to a purple color), and a second chromosome containing the c*2-gfp* allele that makes the seeds yellow (not pigmented) and fluorescent under blue light. Plants of this genotype (ABS4 *C2*/*c2-gfp*) and carrying LexA- CENH3 were crossed as females to a *c2*/*c2* line which has colorless kernels. About 1% of the kernels from this cross are both pigmented and fluorescent, and often contain newly formed neochromosomes. In the original screen, we found four neochromosomes that we named Neo4L-1, Neo4L-2, Neo4L-3, and Neo4L-4 ^21^. To simplify the nomenclature and to accommodate new results, we have renamed the original four as 4b(1), 4b(2), 4b(3), and 4b(4). Using kernels from the same genetic screen, we have now identified and characterized six additional neochromosomes, 4b(5), 4b(6) 4b(7), 4b(8), 4b(9), and 4b(10). In all cases LexA- CENH3 was only used to activate the ABS4 centromeres; after the neochromosomes were identified, we chose individuals for further crossing that lacked LexA-CENH3 (which is unlinked to ABS4).

For all ten neochromosomes, we used fluorescent in situ hybridization (FISH) to confirm that they were present, that they carried ABS4, and to demonstrate that they had acquired telomere repeats at the broken ends. To visualize the location of native centromere 4, we used a probe to a pericentromeric repeat called Cent4 ^29^. We also carried out whole genome Illumina sequencing of the neochromosome lines (Fig. 2, Fig. S1). All ten appeared to contain the entirety of the original sequence to the right of ABS4 (that is, between ABS4 and the telomere of 4L). However, due to the effects of BFB and the random nature of chromosome breakage, each chromosome differs with respect to the sequence to the left of ABS4. The shortest is 4b(6) which appears to have broken right next to ABS4, leaving very little of a short arm on the neochromosome. The longest is 4b(10) which contains most of the sequence between Cent4 and ABS4. Two of the neochromosomes contain evidence of duplications (4b(2) and 4b(10)) that are likely a result of the BFB cycle that preceded the formation of the stable neochromosomes (Fig. S1).

**Figure 2.**
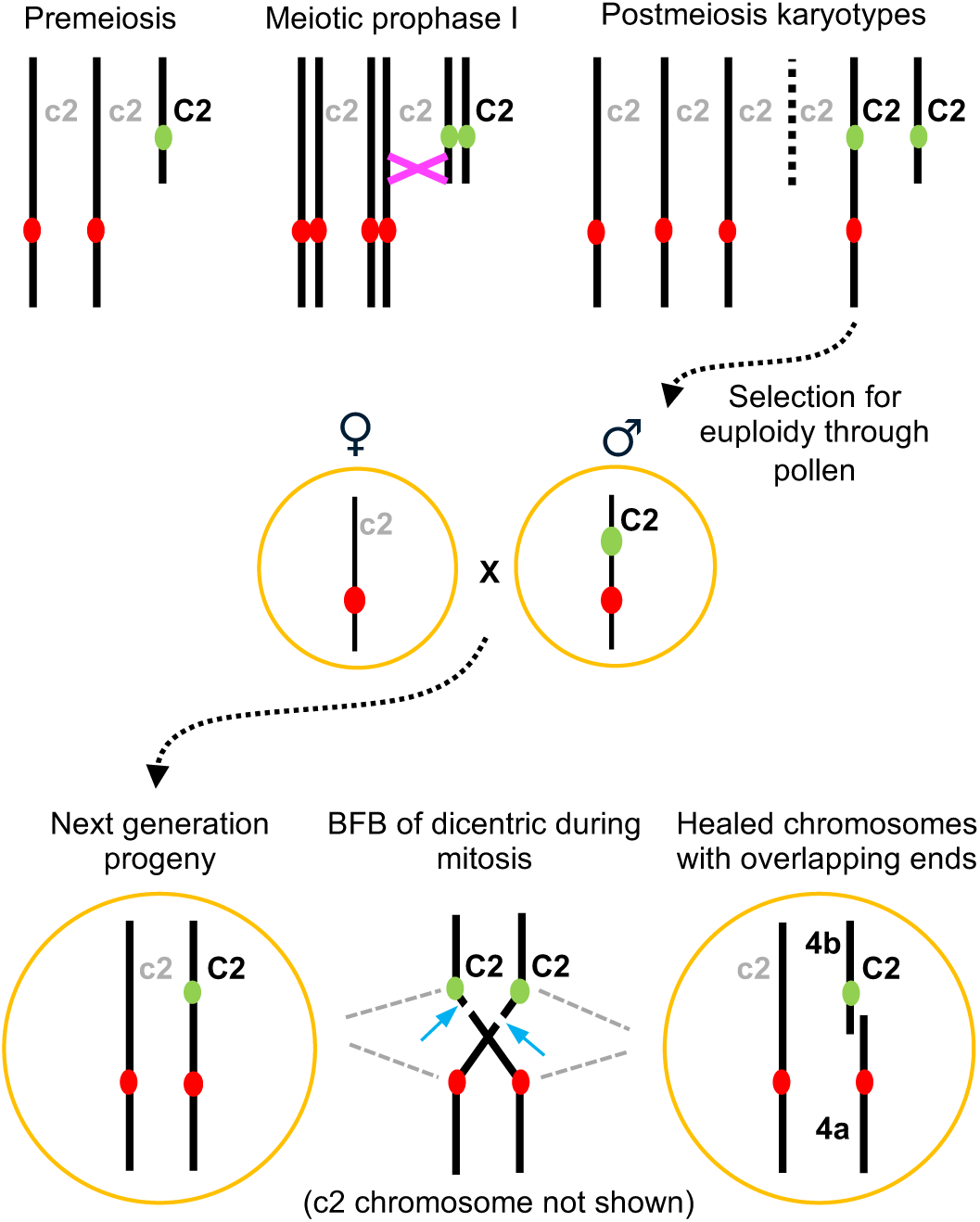
A model for the formation of 4a chromosomes from partially trisomic lines. Trisomic chromosomes pair irregularly but undergo normal recombination. Recombination between normal 4 and 4b (magenta X) can create a dicentric chromosome that is transmissible through pollen (as well as a non-transmissible acentric chromosome (dashed line)). In the next generation, the dicentric chromosome will undergo BFB and two newly broken 4a and 4b chromosomes, often with overlapping ends, can result. The BFB panel shows mitotic anaphase, where thin grey lines indicate spindle microtubules and light blue arrows indicate the positions of new breaks. It is also possible to produce pollen containing 4a and 4b chromosomes with overlapping ends as a direct outcome of meiosis (Fig. S4).

We also carried out CENH3 CUT&Tag to interpret the location and size of the newly formed centromeres. In most cases, CENH3 was localized over the ABS4 insertion as well as flanking sequences to the left and right of ABS4, occupying regions of approximately 1.5 Mb to 2.3 Mb, similar in size to native centromeres ^30–32^. In the flanking genomic sequences, CENH3 is enriched in intergenic spaces and dips to low levels directly over genes similar to cases where genes have been observed in native centromeres (Fig. 1) ^33–35^. There were also smaller, unexpected peaks of CENH3 several Mb away from ABS4 in 4b(1), 4b(5) and 4b(6) (Fig. S2). These peaks may represent cases of spontaneous accumulation of CENH3 that sometimes occur on acentric fragments e.g. ^36^, or structural rearrangements of the 4b chromosomes such as deletions or duplications that bring these apparently distal sequences on the reference in close proximity to the ABS4 array on the neochromosome (we have not assembled the genomes of these neochromosomes).

### Selection for 4a chromosomes

The biology of plant reproduction has a major effect on the inheritance of aneuploid gametes. Both the female and male gametophytes (pollen) are multicellular haploid structures that express genes that support their function. Because pollen tubes compete with each other to reach the egg cell, normal euploid pollen regularly outcompetes aneuploid pollen with altered gene dosage ^37^. In contrast, female gametophytes do not compete with each other and are more tolerant of altered gene dosage. The effect is that extra chromosomes are transmitted at higher frequencies through the female than the male ^38^.

Most of the neochromosomes were recovered as trisomics carrying three alleles, *C2*, *c2-gfp*, and c2, where *C2* was on the neochromosome. When plants of this genotype are crossed as a female, we observed that ∼20-30% of the kernels carried *C2*, and when they were crossed as a male, we observed that about ∼8-12% of the kernels carried *C2* (Table S1). We initially thought that most of the C2 kernels from both female and male crosses were trisomic ^21^. Whether the progeny are indeed trisomic can be estimated by the frequency of *C2* kernels that carry *c2*-*gfp* (both pigmented and fluorescent). Assuming random assortment, about half of the *C2*-carrying trisomic individuals will have *c2*-*gfp* and half will have *c2*. We observed this to be true in female crosses, where on average 51% of the progeny were fluorescent. However this was not true in male crosses, where only 15% carried *c2*-*gfp* (Table S1). These results suggested that most of the *C2* progeny from male crosses were not trisomic and likely did not have the original neochromosome. To test if recombination on the neochromosome might be higher than expected in male crosses, we designed PCR primers over a 26 bp insertion/deletion polymorphism that differentiates the *C2* from *c2* alleles used in our crosses. We found that recombination between *C2* and ABS4 was 14% in female crosses and 55% in male crosses (Table S2; p < 0.01 based on a logistic regression test). It is unlikely that recombination frequencies are elevated in males by three-fold genome wide ^39^. Rather, we believe that recombination rates are similar in male and female, but that recombinants with *C2* but not ABS4 are selectively recovered because they are transmitted in euploid pollen that outcompetes aneuploid pollen. In other words, by choosing *C2* kernels, we biased the sample towards recombinants (Fig. 2).

The realization that trisomic neochromosome progeny are rarely recovered from a male cross led us to reinterpret our 4b(1) data. Unlike all other neochromosomes, which were identified through female crosses, 4b(1) was found fortuitously in the progeny of a male cross ^21^. Our Illumina sequencing of this line seemed to show that the region to the right of ABS4 was disomic, not trisomic as was the case for all others (Fig. 1B). We further noted that the transmission of 4b(1) was consistently higher than what we observed in other lineages, exceeding 40% through female crosses (Table S1). We then analyzed the chromosomes in early-stage prophase cells (where the chromosomes are longer) and noted that Cent4 was present on two clearly different sized chromosomes (Fig. 3F,G). This size difference is apparent in the kernels from the first generation. As confirmation, we crossed 4b(1) (carrying *C2*) to *c2-gfp*, and crossed the resulting *C2*/*c2-gfp* heterozygote to a *c2*/*c2* line. The progeny of these crosses segregated for only pigmented and fluorescent kernels (*C2*/*c2* and *c2-gfp*/*c2*), with no non-pigmented and non-fluorescent kernels nor pigmented and fluorescent kernels (*c2*/*c2* and *C2*/*c2-gfp*), demonstrating the region to the right of ABS4 is disomic. If 4b(1) had been trisomic, the expectation would be segregation of all combinations of the three alleles. These data indicate that two broken chromosomes were recovered together in the original isolate, both 4b(1) and a truncated form of chromosome 4 that we will refer to as 4a(1).

**Figure 3.**
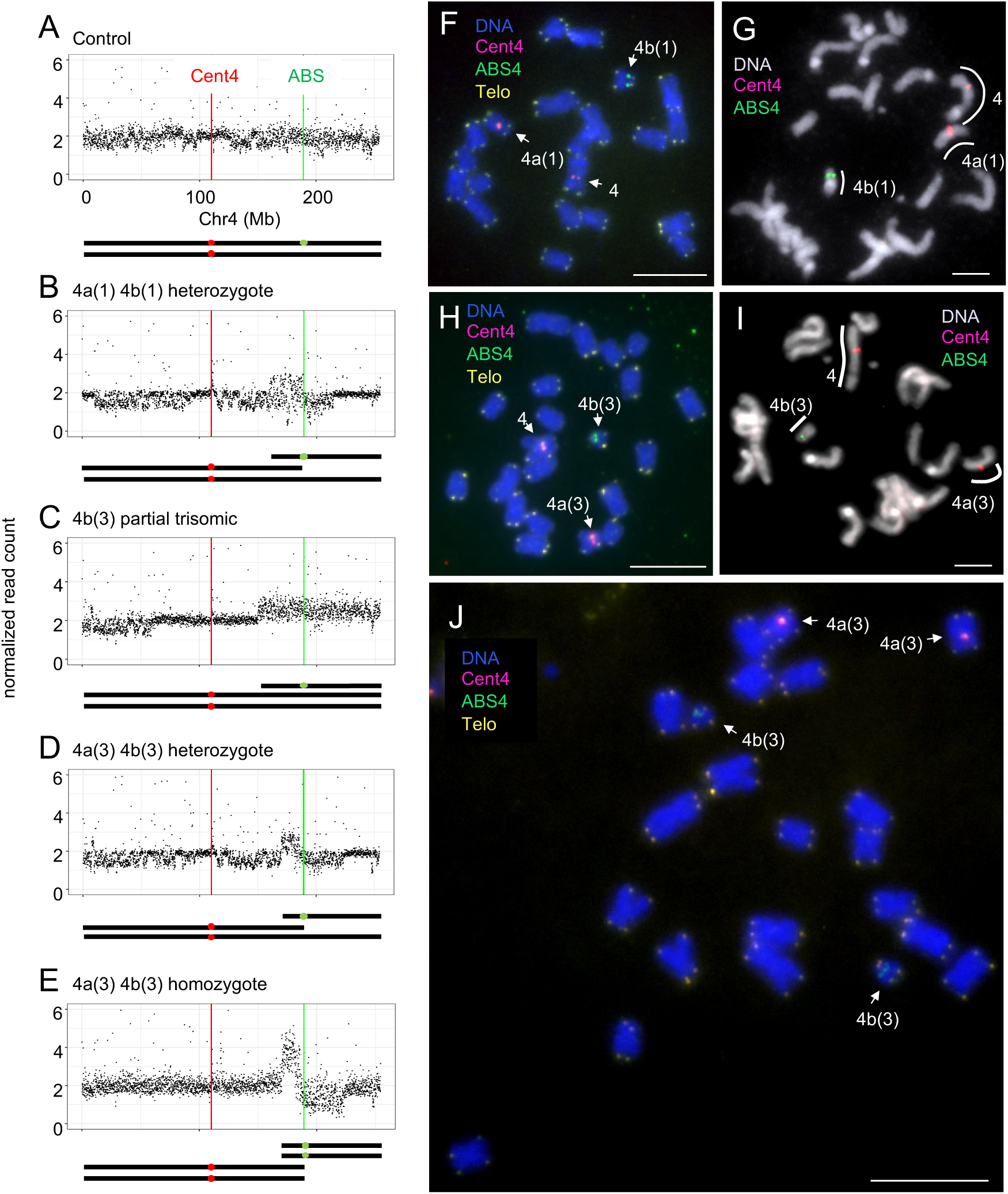
Analysis of 4a 4b lines. **A-E**) Illumina sequencing of plants carrying 4a(1) 4b(1) and 4a(3) 4b(3) pairs. The Y axis indicates the relative ploidy, expressed as normalized read count on 100-Kb intervals. The control is heterozygous for the karyotypically normal ABS4 chromosome and the c2 tester chromosome. The 4a 4b lines have been backcrossed to various degrees and show some areas of homozygosity, evidenced by narrower distributions of read counts. Cartoon representations of the karyotypes are shown below each genotype. **F**) FISH image of a 4a(1) 4b(1) heterozygote in late prophase. **G**) FISH image of a 4a(1) 4b(1) heterozygote in early prophase. **H**) FISH image of a 4a(3) 4b(3) heterozygote in late prophase. **I)** FISH image of a 4a(3) 4b(3) heterozygote in early prophase. **J**) FISH image of a 4a(3) 4b(3) homozygote in late prophase. The Telo label within panels indicates telomeres. Scale bars, 10 μm.

The discovery of the 4a(1)-4b(1) pair led us to reassess the pedigree and transmission data for 4b(3). In the second generation we propagated 4b(3) from a male cross. When one of the progeny from the male cross were planted and crossed again, the transmission was unusually high, showing 47% transmission of *C2* through the female and 35% through the male (Table S1, crosses involving KD4310-2). Progeny from this lineage continued to show female C2 transmission in excess of 40%, suggesting that the 4b(3) chromosome was segregating in a disomic manner and that a 4a(3) chromosome may be present. Analysis of early stage prophase cells demonstrated that this was the case (Fig. 3H,I). Importantly, we had Illumina sequenced the first generation plant, and at this point the line was trisomic and the breakpoint of the 4b chromosome was at position ∼150 Mb on the ABS4 reference genome (Fig. 3C). After crossing as a male and the progeny propagated, there were two new breakpoints: The breakpoint on the 4a chromosome at position ∼182 Mb and breakpoint on a now-shorter 4b chromosome at position ∼170 Mb (Fig. 3D).

These data suggest that when a trisomic 4b line is crossed as a male, a subset of the 4b chromosomes have undergone recombination on the short arm (between ABS4 and Cent4) to reconstitute a dicentric chromosome (Fig. 2). When the newly formed dicentric chromosomes begin mitotic divisions, they are expected to undergo BFB. Supporting this view is the fact that about 8% of the C2 kernels from male crosses have non-pigmented sectors that are suggestive of BFB (See Table S1; though we initially thought sectors reflected centromere instability ^21^). As a test of this hypothesis, we carried out FISH on a sibling of the plant that gave rise to 4b(3). The data show that this lineage contains a different 4b chromosome (with a much shorter arm, appearing to be telocentric) as well as a 4a chromosome (Fig. S3A, see Table S1 for the 4b(3) lineage). We also grew 8 more kernels from the original male cross and analyzed root tip chromosomes by FISH. Only one seedling contained ABS4, and in this case Cent4 and ABS4 were on the same chromosome and nearly juxtaposed, demonstrating that recombination had indeed occurred on the short arm, and that BFB had shortened the distance between Cent4 and ABS4 (Fig. S3B).

Taken together, the genetic data suggest that most of the C2 kernels from male crosses of trisomic 4b lines are derived from euploid recombinants. Many have undergone recombination on the short arm of 4b and are transmitted as dicentrics that, in the next generation, undergo BFB and form 4a-4b chromosome pairs (Fig. 2). Under this model, breakage events occur during mitosis to form a 4a chromosome and a new 4b chromosome with overlapping ends. An alternative mechanism is that breakage to form a 4a chromosome occurs in meiosis II (Fig. S4). While breakage during meiosis is likely to occur, this mechanism preserves the structure of the original 4b chromosome and would not easily explain the origin of the 4a(3)-4b(3) pair. Under both models, breakage in meiosis or breakage during mitosis in the next generation, telomere formation must occur to stabilize the ends of the breaks.

### Phenotype of the 22-chromosome lines

To obtain pure breeding 22-chromosomes lines, we self-crossed plants heterozygous for 4a-4b pairs and normal *c2-gfp* chromosomes. From 4a(1)-4b(1) lines, we were unable to obtain 4a(1)-4b(1) homozygotes, suggesting that there is a mutation(s) and/or deletion(s) in the 4a(1) or 4b(1) chromosome that makes the homozygote inviable. In contrast, 4a(3)-4b(3) homozygotes were readily obtained from self crosses (homozygote in this context refers only to the centromeres and telomeres that define the 4a(3) and 4b(3) chromosomes). As expected, 22 chromosomes were visible in root tips (Fig. 3A). The resulting plants appeared normal, though they varied in vigor and stature as is typical of selfed lines from mixed genetic backgrounds.

To assess the accuracy of meiosis, we analyzed meiotic prophase stages in three plants homozygous for the 4a(3)-4b(3) pair. At pachytene where homologous chromosomes synapse and recombine, we observed 237 cells with normal tight pairing in the centromere regions (that is, the Cent4 and ABS FISH signals appeared as paired dots; we did not attempt to trace the entire length of each chromosome). At diakinesis and diplotene, the chromosomes condense and shorten, making it possible to differentiate the entirety of the two chromosomes. All 87 cells showed the homologous 4a(3) and homologous 4b(3) chromosomes paired separately and associated by what appeared to be chiasmata (Table S3, Fig. 4A). We observed very few cells in the later stages of meiosis but noted no obvious errors. As a second measure of the accuracy of meiosis, we collected fresh pollen and stained the grains with a vital stain. The data indicate that pollen from homozygous 4a(3)-4b(3) plants was morphologically indistinguishable from control plants (Fig. 4B, Fig. S5). Taken together, the data suggest that plants homozygous for the 4a(3)-4b(3) pair can proceed through meiosis in a normal manner.

**Figure 4.**
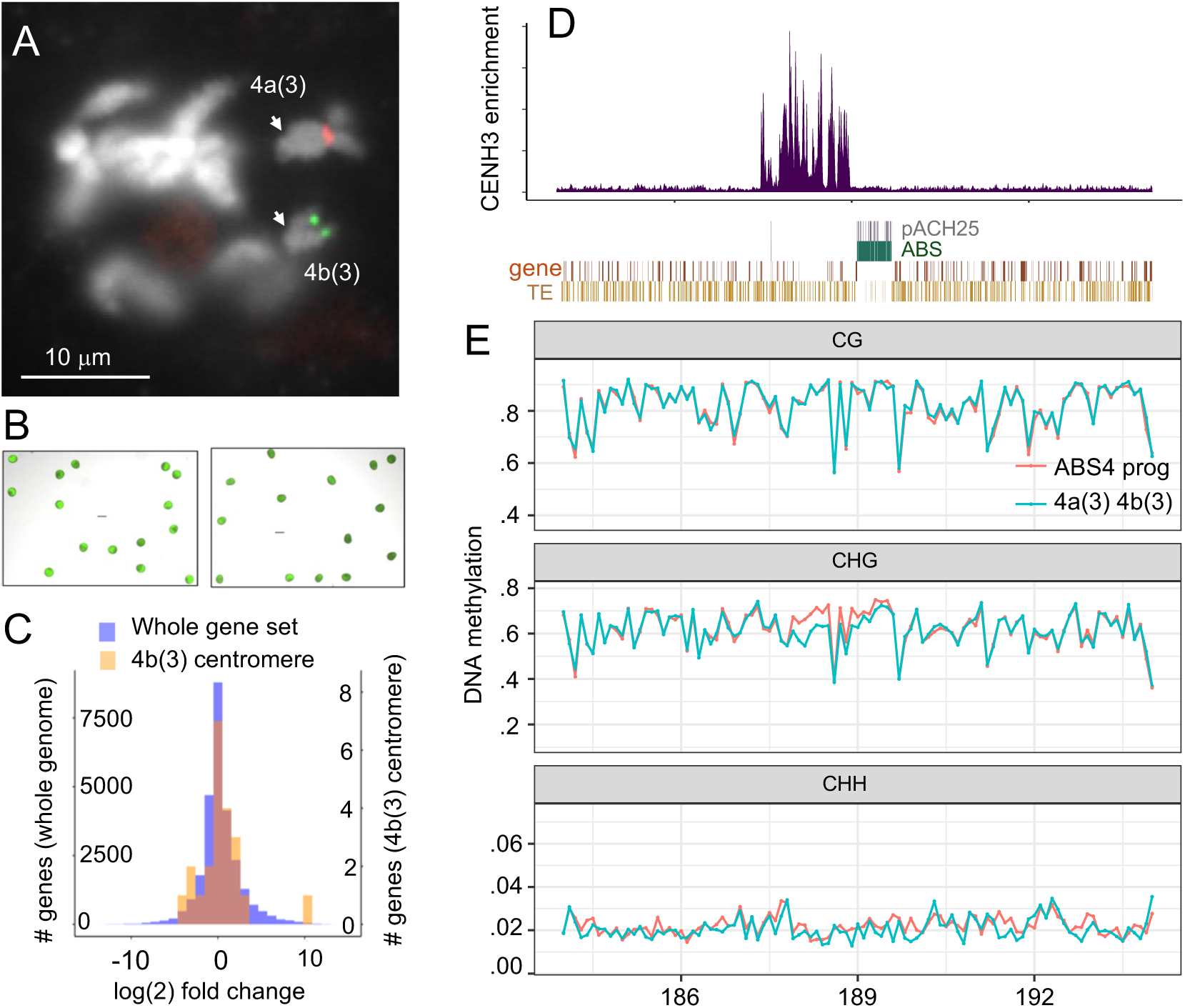
Phenotypes of homozygous 4a(3) 4b(3) lines. **A**) Meiotic cell in late diplotene showing the pairing of the homozygous 4a(3) and 4b(3) chromosomes. Arrows point to chiasmata. DNA is in white, Cent4 in red, and ABS4 in green. **B**) Pollen in homozygous 4a(3) and 4b(3) plant (left) and normal karyotype c2/c2 line (right) stained with fluorescein diacetate. Scale bars (suspended in the images) are 100 μm. **C**) Relative gene expression between 4a(3) 4b(3) homozygotes and the ABS4 progenitor line. One gene (Zm00056aa025500) shows notably higher expression in the 4a(3) 4b(3) line than in the ABS4 progenitor. Figure S7 shows expression of all 22 genes in the four replicates of each genotype. **D**) CUT&Tag profile in the homozygous 4a(3) 4b(3) line and annotation features in the local area. **E**) DNA methylation profiles in each of the CG, CHG, and CHG contexts in the same plants used for RNA expression analysis in C.

We obtained seven full ears from self- or sib-crossed homozygous 4a(3)-4b(3) lines (Table S4, Fig. S6). We also made both female and male crosses to a line that is homozygous for both the recessive c2 allele and an allele of the *R1* gene (*R1-scm2*) that promotes strong anthocyanin formation in the scutellum (a visible part of the seed derived from the embryo). These crosses were designed to assess whether 4b(3) might be subject to centromere erasure and chromosome loss in the embryo, similar to the centromere mediated haploid induction process ^40,41^. Of 432 kernels from female crosses and 889 kernels from male crosses, we observed no kernels with colorless scutella (Table S4). We did, however, observe a low frequency of sectored kernels in male crosses (1%). Sectored kernels likely represent cases of recombination between the short overlapping regions of 4a(3) and 4b(3), leading to dicentric chromosomes and BFB.

### Effect of the 4b(3) centromere on gene expression and DNA methylation

Having 4b(3) in a homozygous state allowed us to assess DNA methylation and gene expression in the CENH3-occupied area. To this end, we first performed CUT&Tag on the homozygous line and observed that in the three generations since the first CUT&Tag was carried out ^21^, the centromere location had shifted entirely into genomic sequence to the left of ABS4 and occupied a region of ∼1.5 Mb. While centromere shifting is not unexpected ^30^, the new position is fortuitous because short reads can be more accurately mapped to these genomic sequences.

Prior analyses of native centromeres have shown that CENH3-enriched regions and pericentromeric regions have high levels of DNA methylation in the CG context, while the CENH3-enriched regions have somewhat reduced CHG methylation ^6,42^. To test if the same trends were observed in the 4b(3) centromere, we grew four plants from the ABS4 inbred (no centromere at ABS4) and four plants that were homozygous for the 4a(3)-4b(3) pair and carried out Enzymatic Methyl sequencing (EM-seq) of leaves. Analysis of the methylation patterns revealed no visible changes in CG methylation throughout the ABS4 region (Fig. 4E, Fig. S7). However, consistent with prior data from native centromeres, there was a slight reduction in CHG methylation in the CENH3-enriched area (from ∼70% to 60%). DNA methylation within and around genes was largely unaffected by the formation of the 4b(3) centromere.

When a centromere shifts into genomic regions, the expression of genes in the affected area might be altered. To test this, we used tissue from the same eight plants (four ABS4 control and four 4a(3)-b(3)) for comparative RNA-seq analysis. Overall, 8,222 of 23,204 genes with detectable expression were differentially expressed (Wald test, Bonferroni adjusted p-value < 0.01). A total of 4,493 genes showed increased expression in the homozygous 4a(3)-4b(3) plants and 3,729 genes showed decreased expression relative to the ABS4 control. Of the 22 genes in the CENH3 enriched region with detectable expression, four had increased expression in 4a(3)-4b(3) plants, and one had decreased expression (Fig. 4C, Fig. S7). The proportion of genes with increased or decreased expression in the CENH3 enriched region was not significantly different from the whole-genome average (Chi-squared test p-value = 0.2071 for increased, 1 for decreased). However, one gene, which is nearly inactive in the progenitor ABS4 line, was sharply up-regulated (∼10-fold) in the 4a(3)-4b(3) line, perhaps suggesting that CENH3 can in some cases alter cis-regulatory features of nearby genes.

## DISCUSSION

Methods for routine centromere engineering could be used to dramatically alter karyotypes, restructure recombination landscapes and speed the development of artificial chromosomes ^1,43,44^. In this study we describe a pathway for the formation of stable new karyotypes from lines with newly formed centromeres. The key criterion is that the new centromere be on the same chromosome as the original centromere and far enough away that meiotic recombination between the two centromeres is possible. When starting with a partially trisomic line, recombinants of this type are favored when the plants are crossed as a male (Fig. 2). The dicentric chromosomes formed by recombination undergo BFB and can be converted to chromosome pairs with overlapping ends, which as we show here can be transmitted to subsequent progeny and recovered as new stable karyotypes.

The rapid formation of telomeres on newly broken ends is critical for the formation of new chromosomes. McClintock demonstrated that BFB cycles usually terminate early in development by a process of healing (de novo telomere formation) ^26,27^. Later studies in wheat showed that dicentric chromosomes underwent BFB until about 4 weeks after germination, at which point most of the broken ends were healed with telomeres visible by in situ hybridization^45^. Four telomere addition sites were sequenced, and the sites of addition had 2-4 nucleotides of homology to the telomere repeat ^46^, consistent with biochemical studies showing that plant telomerases can initiate new telomeres on a variety of templates ^28^. In our work we did not visualize telomeres until the second (or later) generation after the neochromosomes were formed, and in all cases telomeres were visible by in situ hybridization on the recently broken ends.

The ten 4b centromeres described here cover a minimum of 1.5 Mb of sequence, similar to native centromeres ^47^, and in all cases CENH3 spread into the flanking genic areas. For the homozygous 4a(3)-4b(3) line, we found no evidence for reduced expression within the CENH3 domain relative to the progenitor ABS4 chromosome beyond normal gene expression variation. The fact that centromeres can form among genes has also also been documented in native centromeres and spontaneous neocentromeres in both plants and animals ^33,36,48,49^. The forces that drive and limit CENH3 spreading, and the mechanisms that exclude CENH3 from genes are not yet understood. Newly formed centromeres also lack the long spans of flanking pericentromeric heterochromatin that typically surround the centromere cores of native centromeres. On the 4b(3) centromere we observed no increase in DNA methylation that might indicate an elevated or altered heterochromatin environment (Fig.4E). These data support the view that pericentric heterochromatin is the result of a gradual build up of repetitive sequences in low-recombination regions within and surrounding centromeres, and is not required for centromere function ^49–52^.

Perhaps the most compelling case that the new telomeres and the 4b(3) centromere are functional, and not impairing gene expression or other essential processes, comes from the fact that the 22-chromosome maize lines grew and reproduced as well as any standard laboratory strain. However, despite its normal phenotype, the 22-chromosome line is no longer compatible with breeding programs involving standard (20-chromosome) maize lines. Whenever a 22 chromosome line is crossed to a standard maize line, recombination between the two centromeres on chromosome 4 will lead to dicentric chromosomes and further structural rearrangements. For this reason, major karyotype alterations are thought to be one form of genetic isolation that leads to speciation ^53^.

The fact that 22-chromosome maize lines are not compatible with standard maize breeding lines could be seen as an advantage. It may be possible to initiate new breeding programs based on maize lines with altered karyotypes. In this way proprietary maize lines could be durably marked by their unique karyotypes. Engineered gene clusters could be placed on (for instance) the 4b chromosome, perhaps close to the centromere, to ensure that they are not inadvertently introduced into other standard backgrounds. Further, the fission of one chromosome into two separate chromosomes should dramatically alter the recombination landscape, perhaps releasing genetic variation that has the potential to improve maize in new and unpredictable ways ^1^.

## METHODS

### Plant materials

The ABS4 inbred was developed by self-crossing the original transgenic line in a Hi-II genetic background five times ^23^. The *c2-gfp* allele was obtained as stock tdsgR64H07 and the *c2 R1-scm2* strain as stock X25B from the Maize Genetics Cooperation Stock Center in Urbana, IL. Plants were grown and crossed in the UGA Botany greenhouses.

### Genotyping

The presence of the ABS4 insertion was assayed using primers abs4-p3 (5’- TCCTCCGGAGTACCGTCT-3’) and abs4-p4 (5’-AGCCAGGCGGATAGAAGC-3’) whereas the wild-type locus was amplified using primers abs4-p1 (TACCCTGGTTAGAGGGAGCC) and abs4-p4 ^21^. To test for the presence of the dominant C2 allele by PCR, we developed primers (C2-F1, 5’-CACAGCGTCCCCATCACC-3’ and C2-R1 5’-GAACGAGACGACGACGAATTG-3’) that amplify a sequence over 26 bp insertion-deletion polymorphism in the 3’ UTR of the c2 gene (the *C2* allele in the ABS4 inbred has the insertion, whereas our standard c2/c2 line has the deletion). Fluorescence from the *c2-gfp* allele was visualized with blue light source and orange filter (Clare Chemical HL34). In cases where both the C2 allele and c2-gfp allele were present, it was often necessary to sand off (with sandpaper) a small portion of the aleurone to visualize the fluorescence.

### HiFi sequencing and assembly of the ABS4 inbred

The ABS4 homozygous line in the Hi-II background was grown in the greenhouse of the University of Georgia, Athens, and the two top leaves of a mature plant were sent to the Arizona Genomics Institute (AGI) in Tucson, Arizona. DNA was extracted using the CTAB method ^54^, and library preparation was performed with the SMARTbell Express Template Prep Kit 3.0. The final library preparation included size selection for fragments ranging from 10–25 kb using the Blue Pippin system (Sage Science). Sequencing was conducted on a PacBio Revio system in Circular Consensus Sequence (CCS) mode. The read length N50 was determined to be 22,840 bp. Genome assembly was carried out using Hifiasm v0.19.4 ^55^ with the ‘-t 256’ and ‘-l 0’ parameters to enable phasing, sequence error correction, and assembly. The optional parameter -u was initially tested but was disabled due to increased assembly errors. The final assembly produced 1,136 contigs with a total length of ∼2.3 Gb and a contig N50 of 34,924,731 bp.

Ragtag v2.1.0 ^56^ was used to scaffold the contigs using the A188 reference genome ^24^. The sequenced ABS4 genome exhibited a low level of heterozygosity, particularly on chromosome five, where heterozygous contigs were shorter and had lower coverage. Ragtag v2.1.0 does not distinguish heterozygous contigs and can place genome duplicates into the final reference. A customized algorithm was developed to identify and filter duplicate contigs by calculating the percentage overlap between consecutive contigs using Ragtag v2.1.0-generated PAF and AGP files. Contigs with over 99% overlap with the preceding contig were removed. Contig ptg000071l was misassembled due to a repetitive area consisting of TAC trinucleotide tandem repeats. The RagTag v2.1.0 ‘correct’ command was used to split this contig and re-scaffold with the scaffold command.

We identified four primary contigs containing ABS and pACH25 sequences. Two of these contigs were placed by Ragtag, as they contained maize genome sequences, while the other two unplaced contigs were found to have genomic regions with 100% identity to the placed contigs. We manually scaffolded these two unplaced contigs into the final assembly by joining the overlapping regions. To plot the synteny between the A188 and the ABS4 genome, syntenic genes were paired by gene name and the data visualized using KaryoploteR ^57^.

### Annotation of the ABS4 inbred genome

Gene annotation was performed with Liftoff v.1.6.3 ^58^ using the A188 reference and gene annotation ^24^ as input. Transposable elements were annotated using EDTA v2.1.0 with ‘ -- species Maize --step anno --overwrite 1 --cds --u 1.3e-8 --threads 18’ parameters ^59^.

To estimate the number of LexO sites in the ABS4 array, the large (second) insertion region was extracted using BEDTools v2.31.0 ^60^ with the ‘getfasta’ command and a nucleotide BLAST database was built using the ‘makeblastdb’ command. The LexO motif (TACTGTATATATATACAGTA) was queried against the database using the ‘blastn-short’ command and only exact matches to all 20 bp counted. By this measure the estimated number of LexO binding sites is 2,783.

### Fluorescent in situ hybridization of mitotic and meiotic chromosomes

The preparation of metaphase spreads from root tip chromosomes and subsequent fluorescent in situ hybridization (FISH) was carried out as previously described ^21^, using FITC-labeled oligos for ABS, Cy5-labeled oligos for Cent4, and a Cy3-labeled oligo (5’- TTTAGGGTTTAGGGTTTAGGG-3’) for telomere repeats. The preparation of meiotic samples and subsequent FISH was carried out as previously described ^21^ using FITC-labeled oligos for ABS and Cy5-labeled oligos for Cent4. Cells were imaged using a Zeiss Axio Imager.M1 fluorescence microscope with 63X or 100X plan-apo Chromat oil objectives. Data were analysed using Slidebook software (Intelligent Imaging Innovations).

### Whole genome Illumina sequencing and analysis

Whole genome sequencing libraries were prepared with KAPA HyperPrep Kits (KK8502) using 50 ng sonicated input DNA with NEXTflex® DNA Barcodes (Bioo Scientific NOVA- 520996), reducing all reaction volumes by ½, and amplifying with 4 or 5 cycles of PCR using the NEXTFlex primer mix. Libraries were sequenced paired-end with 150-nt reads. Adapter sequences were trimmed using Trim Galore v0.6.7 ^61^ with the --fastqc, --gzip, and --paired -a AGATCGGAAGAGC parameters. Trimmed reads were then aligned to the reference genome using BWA-mem v0.11.3 with the -M parameter ^62^. Three reference genomes—ABS4 assembly, Mo17, and W22—were tested, with the W22 genome ^63^ yielding the most consistent results. A 100-kb window file for the W22 genome was generated using BEDTools v2.31.0 ^60^ with the makewindows tool. Then the intersect tool with ‘-c’ was employed to calculate read counts for each window. The output file was processed in R, where the normalized read count was calculated as the read count of each 100-kb window divided by the total mapped read count times the total number of 100-Kb windows times 2.

### CENH3 CUT&Tag assays and analysis

Unfertilized ears, varying from 3 to 8 cm in length were used for CUT&Tag with CENH3 antibodies ^64^. Nuclei were purified using a sucrose/percoll cushion method ^65^ and intact nuclei containing 350 ng of DNA used as input for each CUT&Tag reaction with Active motif CUT&Tag-IT kits and amplified with 15 cycles of PCR as described previously ^21^. The nuclei purification method was modified slightly in that two layers of miracloth were used instead of four, nuclei suspensions were filtered only once through cell strainers instead of twice, and 100 micron and 40 micron cell strainers were used instead of 20 micron and 10 micron cell strainers. CUT&Tag libraries were Illumina sequenced paired-end 150-nt. Adapter sequences were trimmed using Trim Galore v0.6.7 ‘--fastqc, --gzip, --paired -a CTGTCTCTTATACACATCT ^61^. Trimmed reads were aligned to the ABS4 assembly using BWA-mem v0.11.3 with the ‘-M’ parameter ^62^. A 10-kb window file for ABS4 assembly was generated using BEDTools v2.31.0 ^60^ with the makewindows tool. Then the intersect tool with ‘-c’ was employed to calculate read counts for each window. The results were visualized using ggplot2 with the geom_area function plotting the CENH3 enrichment peaks and the geom_rect function plotting different genetic elements (genes, TEs, ABS4, and pACH25).

### Testing for elevated recombination between ABS4 and C2 in male crosses

Recombination events, defined as the presence of the *C2* allele without ABS4 were scored by PCR of germinated seedlings. We excluded two samples from the counts that were missing both ABS4 and C2. To examine the relationship between direction of cross and recombination at the ABS4 and C2 loci, we conducted a logistic regression with a custom R script with the model fit Recombination ∼ Cross Direction. Logistic regression showed that the log odds of recombination increased by 1.9994 (95% CI [1.3374, 2.6566], p = 0.002325) in male crosses compared to female crosses.

### Pollen morphology

Freshly released pollen from three plants of each genotype was immersed in an aqueous solution of 5 ng/ml FDA (fluorescein diacetate, Chem Cruz #Sc-294598) in 1.4 M sucrose. FDA assay tests for the integrity of plasmalemma of the vegetative cell, which is an indirect measure of cell viability ^66^. At least 100 pollen grains were counted per plant using images taken with a Nikon Eclipse Ti2 inverted microscope, and any that had partially collapsed cytoplasm or irregular shape were counted as abnormal.

### Gene expression assays

Strand-specific RNA sequencing libraries were prepared from leaves from four individual plants from the ABS4 inbred and four individual plants that were homozygous for 4a(3) and 4b(3) in a complex genetic background including Hi-II in the ABS4 region using the NEBNext Ultra II Directional RNA Library Prep Kit for Illumina following manufacturer’s instructions. Briefly, the RNAs were fragmented for 8 minutes at 94 °C. First strand and second strand cDNA were subsequently synthesized. The second strand of cDNA was marked by incorporating dUTP during the synthesis. cDNA fragments were adenylated at 3’ends, and an indexed adapter was ligated to cDNA fragments. Limited cycle PCR was used for library enrichment. The incorporated dUTP in second strand cDNA quenched second strand amplification, which preserves strand specificity. Libraries were Illumina sequenced paired-end 150-nt. Adapter sequences are AGATCGGAAGAGCACACGTCTGAACTCCAGTCA (read 1) and AGATCGGAAGAGCGTCGTGTAGGGAAAGAGTGT (read 2).

After trimming the adapter with the default parameter with TrimGalore v.6.7, mRNA reads were mapped to the ABS4 inbred using STAR v.2.7 with ‘--outSAMunmapped Within, -- outSAMattributes Standard’ command ^67^. Cufflinks v.2.2.1 ^68^ transformed the Liftoff generated gene annotation gff file to gtf format. Featurecount v.1.6 ^69^ was used to calculate reads count for each exon. ∼89% of reads mapped to the genome but only ∼27% of reads assigned to exons, presumably due to inaccurate annotations generated by Liftoff. Featurecount output files were then channeled into R to calculate the read count for each gene. The DEseq function of Package DESeq2 ^70^ was used to perform the fold changes calculation, relative log expression (RLE), and differential expression gene (DEG) analyses. Of the 26 genes in the 1.5-Mb region of CENH3 enrichment, four had read counts of zero, and were excluded from analysis.

### DNA methylation analyses

Enzymatic Methyl sequencing (EM-seq) libraries were prepared from the same leaves that were used for RNA-seq using a NEBNext Enzymatic Methyl-seq Kit (New England Biolabs #E7120S). The input for each library consisted of 200 ng of DNA that had been combined with 1 pg of control pUC19 DNA and 20 pg of control lambda DNA and sonicated to fragments averaging ∼600 bp in length using a Diagenode Bioruptor. The protocol for large insert libraries was followed with formamide as the denaturing agent, and libraries were amplified with four PCR cycles and Illumina sequenced using paired-end 150 nt reads. EM-seq reads were trimmed of adapter sequence using cutadapt ^71^, parameters -q 20 -a AGATCGGAAGAGC -A AGATCGGAAGAGC -O. Reads were aligned to each genome and methylation values were called using BS-Seeker2 v.2.1.5, parameters -m 1 –aligner=bowtie2 -X 1000 ^72^. The resulting files in CGmap format were processed using CGmapTools v.1.2 ^73^. The replicates of ABS4 homozygous line and 4a(3) 4b(3) homozygous line were merged two by two using the merge2 tool and the 100kb window methylation calculation done with the mbin tool ‘ -c 1, -B 100000’, and the results plotted with gpplot2 wrapped in tidyverse.

## Data Availability

All sequencing data generated in this study have been submitted to the NCBI BioProject database (https://www.ncbi.nlm.nih.gov/bioproject/) under accession number PRJNA874319. The ABS4 reference genome and annotation are available at https://zenodo.org/records/14501890. All code is available at https://github.com/dawelab/Centromere_Engineering.

## ACKNOWLEDGEMENTS

We thank Dong won Kim for technical contributions throughout this study, including genotyping, CUT&Tag assays and library preparation. We thank Meghan Brady for statistical help with recombination data. This study was supported in part by resources and technical expertise from the Georgia Advanced Computing Resource Center, as well as a grant from the National Science Foundation (IOS-2040218).

## SUPPLEMENTAL FIGURES

**Figure S1.**
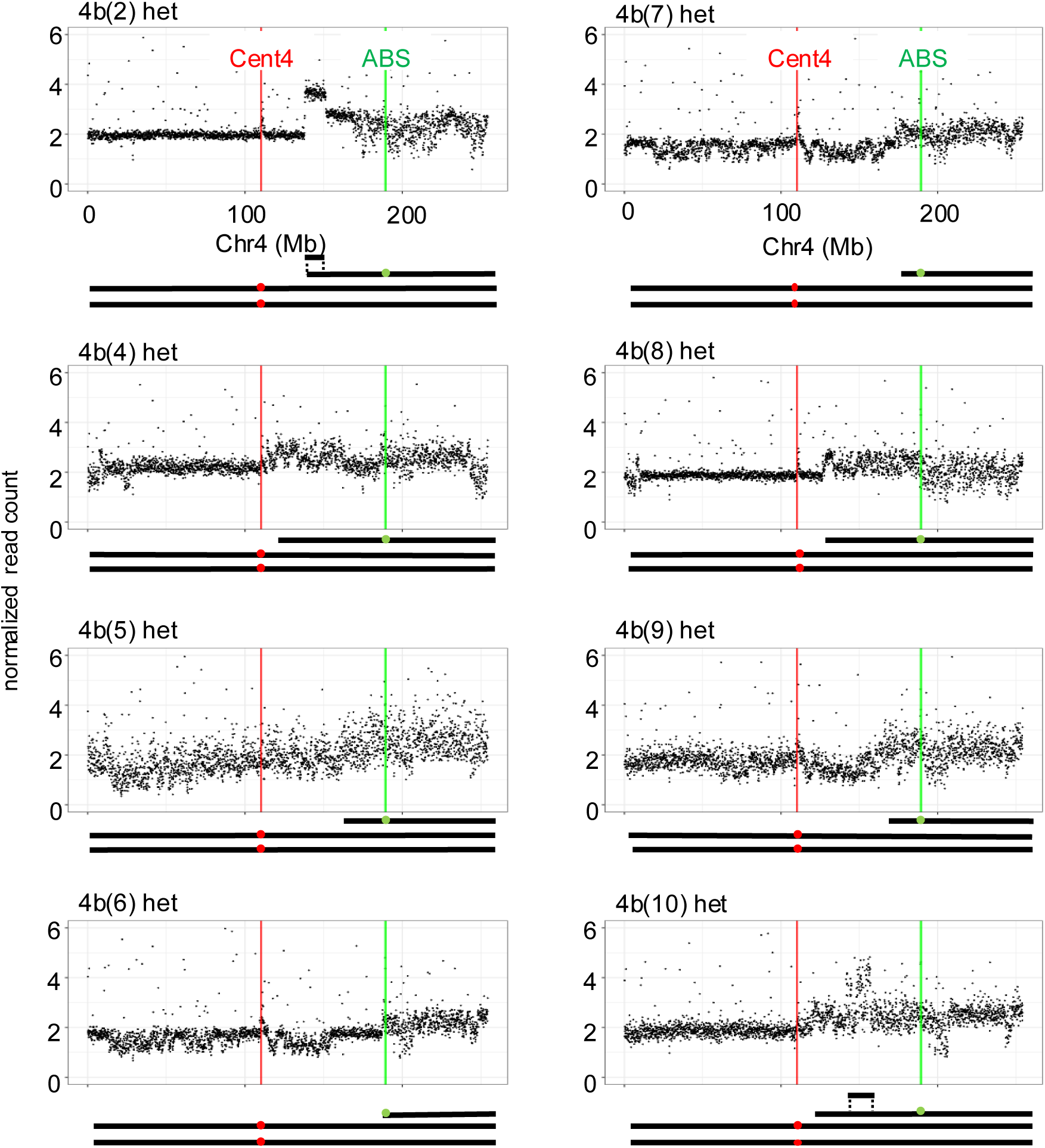
Illumina sequence analysis of eight 4b chromosomes. The Y axis indicates the relative ploidy, expressed as normalized read count on 100-Kb intervals. Cartoon representations of the karyotypes are shown below each genotype. Note that 4b(2) and 4b(10) appear to have duplications of the type expected to result from the BFB cycle. For the control line, 4b(1), and 4b(3), see Fig. 2.

**Figure S2.**
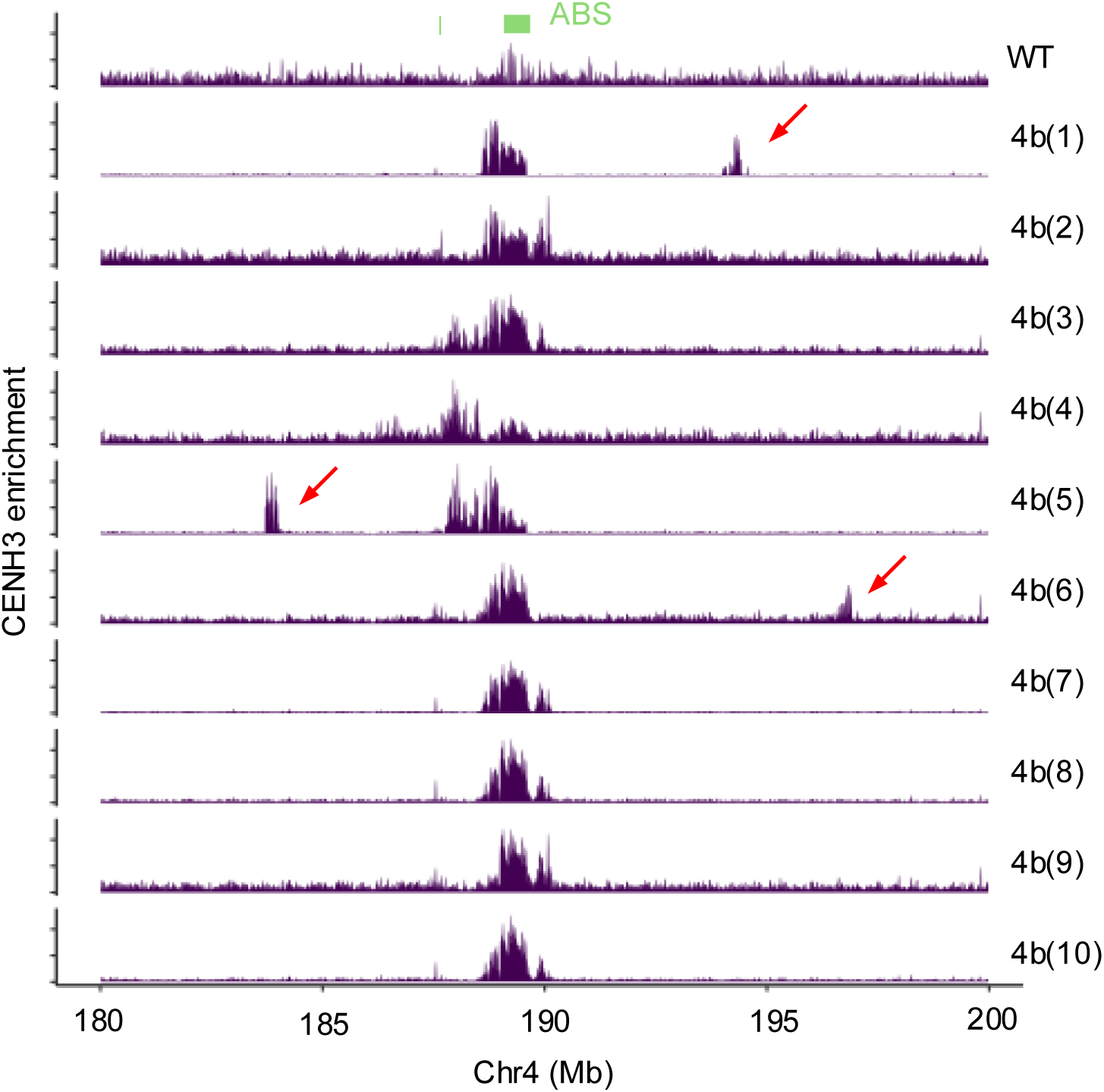
Secondary CENH3 enrichment peaks on neochromosomes. This graph shows a zoomed-out view of the same data displayed in Fig. 1, where only the main CENH3 peaks were shown. Smaller secondary peaks are indicated with arrows.

**Figure S3.**
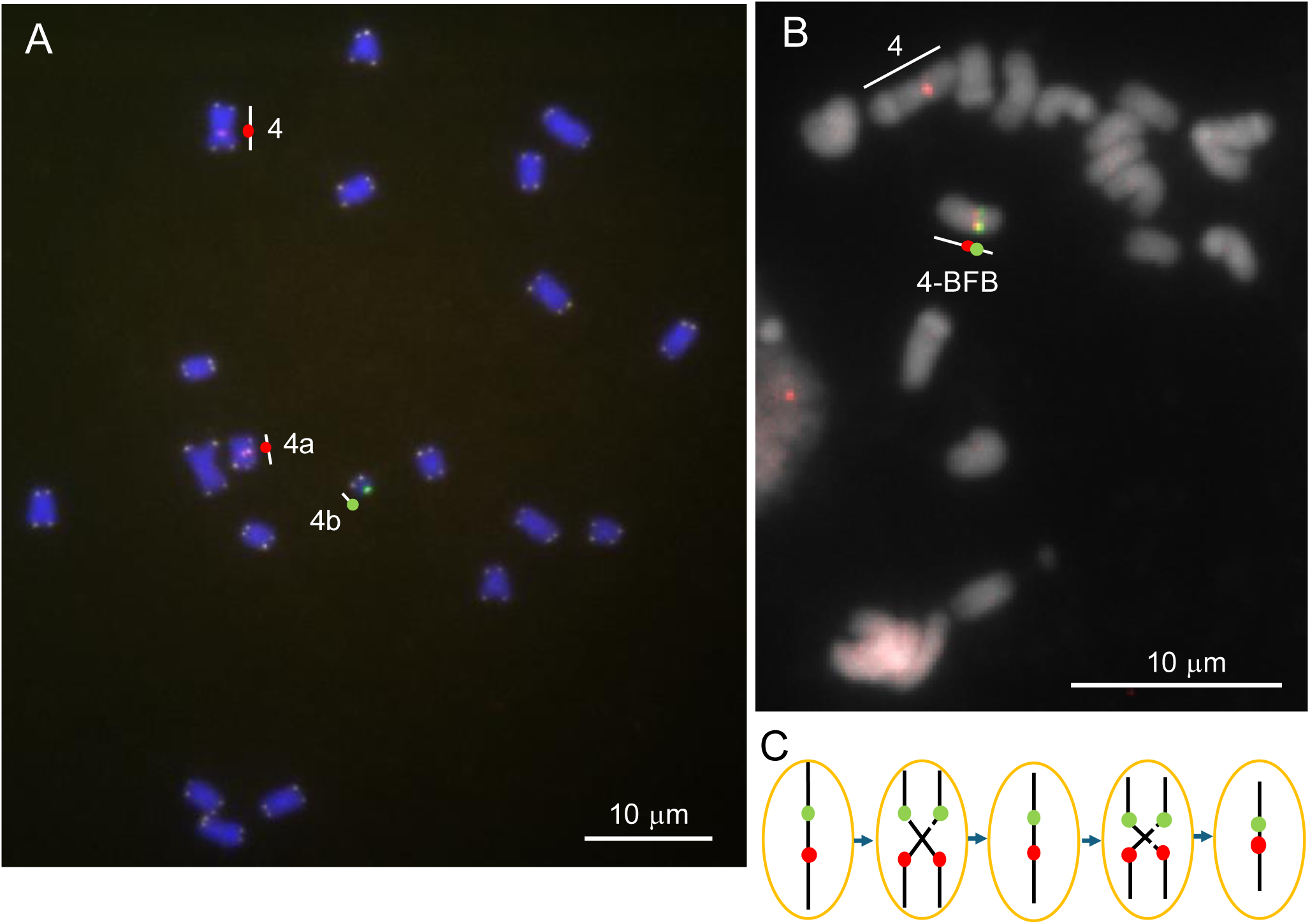
Additional evidence that dicentric chromosomes are transmitted through male crosses of partial trisomic lines. **A**) Root tip FISH from a lineage that originated with the same male cross that led to the discovery of 4b(3) (root here was from KD4310-1 × 4307, see Table S1). This line carries a 4a chromosome and a form of 4b(3) that is telocentric. DNA is in blue, Cent4 in red, ABS in green, and telomeres in yellow. **B**) Root tip FISH from another individual from the male cross that led to the discovery of 4b(3) (seed was from KD4277-2 x KD4303-2; this plant was never grown to maturity). Note that Cent4 and ABS are on the same chromosome, but nearly juxtaposed due to repeated breakage and rejoining during BFB. DNA is in white, Cent4 in red, and ABS in green. **C**) Illustration of how BFB is expected to reduce the distance between two centromeres in a dicentric chromosome.

**Figure S4.**
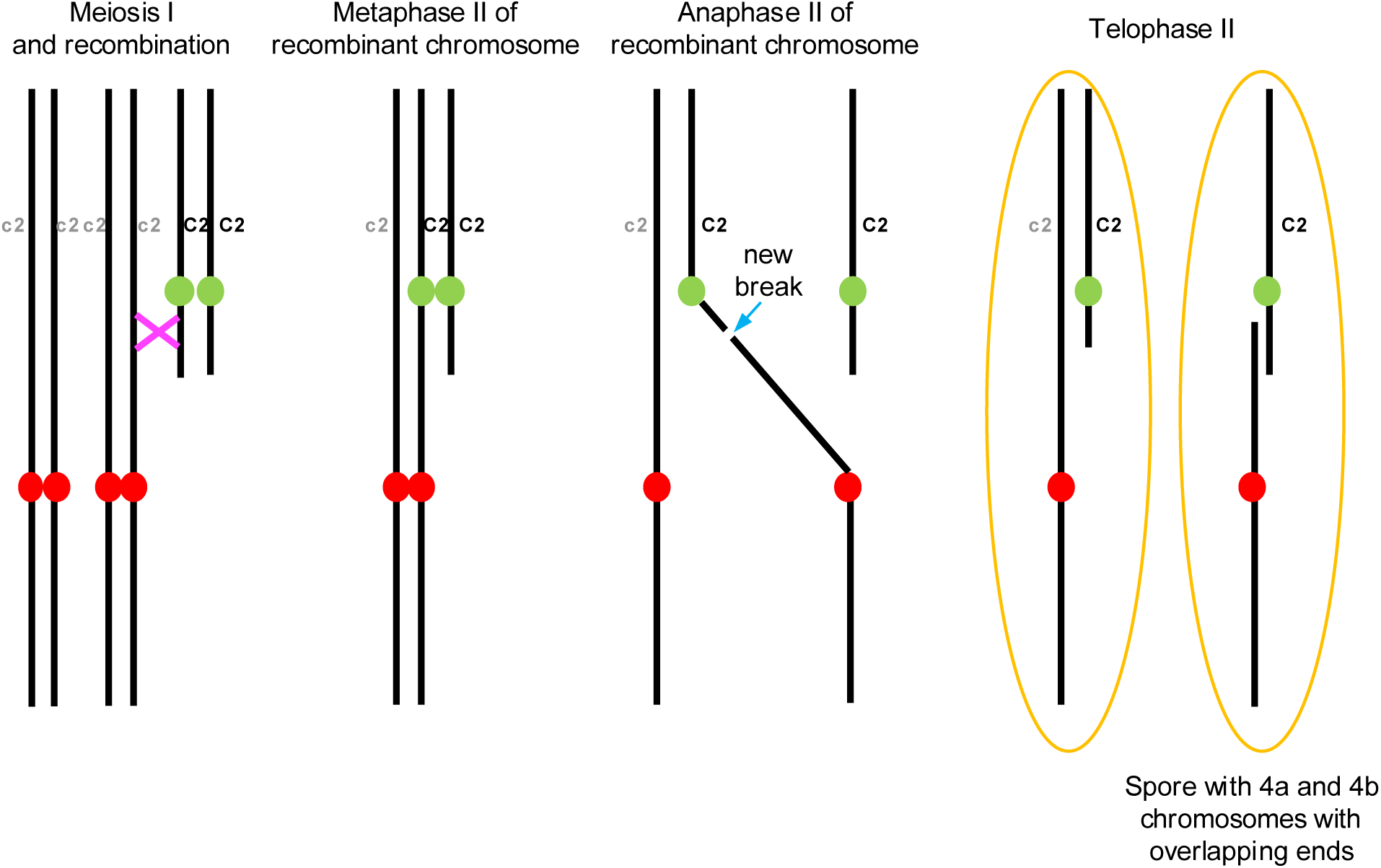
Illustration of how recombination in a partially trisomic 4b line can lead to a meiotic product with 4a and 4b chromosomes that have overlapping ends. This mechanism preserves the original 4b chromosome while creating a 4a chromosome. It only works if the line is partially trisomic. It cannot readily explain the fact that the original 4b(3) chromosome (Fig. 3C) had a longer short arm than the 4b(3) chromosome that was recovered along with 4a(3) (Fig. 3D).

**Figure S5.**
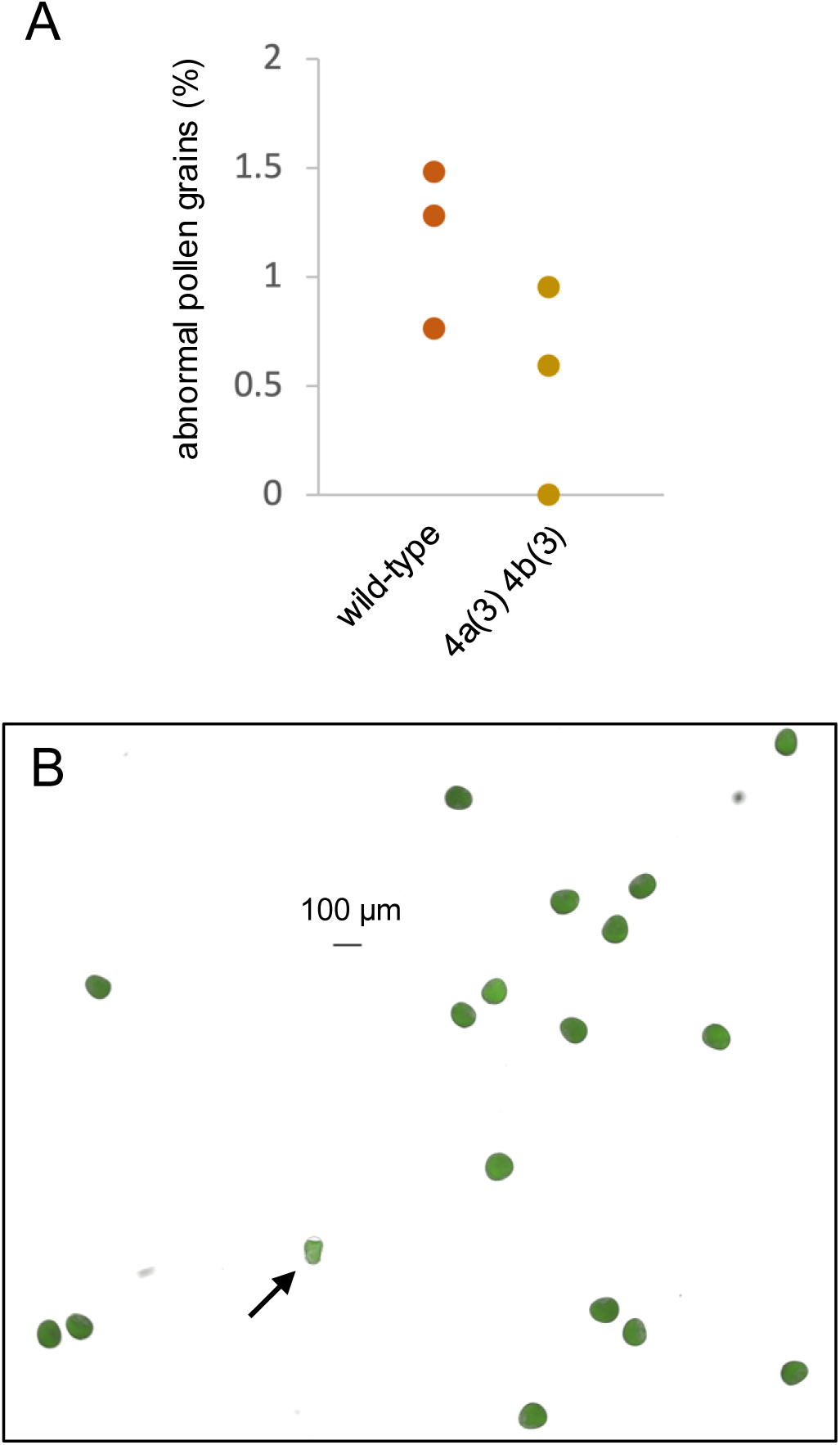
4a(3) 4b(3) homozygous plants produce morphologically normal pollen. **A**) Percentage of abnormal pollen grains from wild-type and 4a(3) 4b(3) homozygous plants. Freshly released pollen from three wild-type (c2/c2 tester line) and three 4a(3) 4b(3) homozygous plants was stained with fluorescein diacetate, imaged, and the frequencies of abnormal pollen grains counted. Each dot indicates the average from one plant. At least 100 pollen grains were counted for each plant. **B**) Example abnormal pollen grain, indicated by arrow.

**Figure S6.**
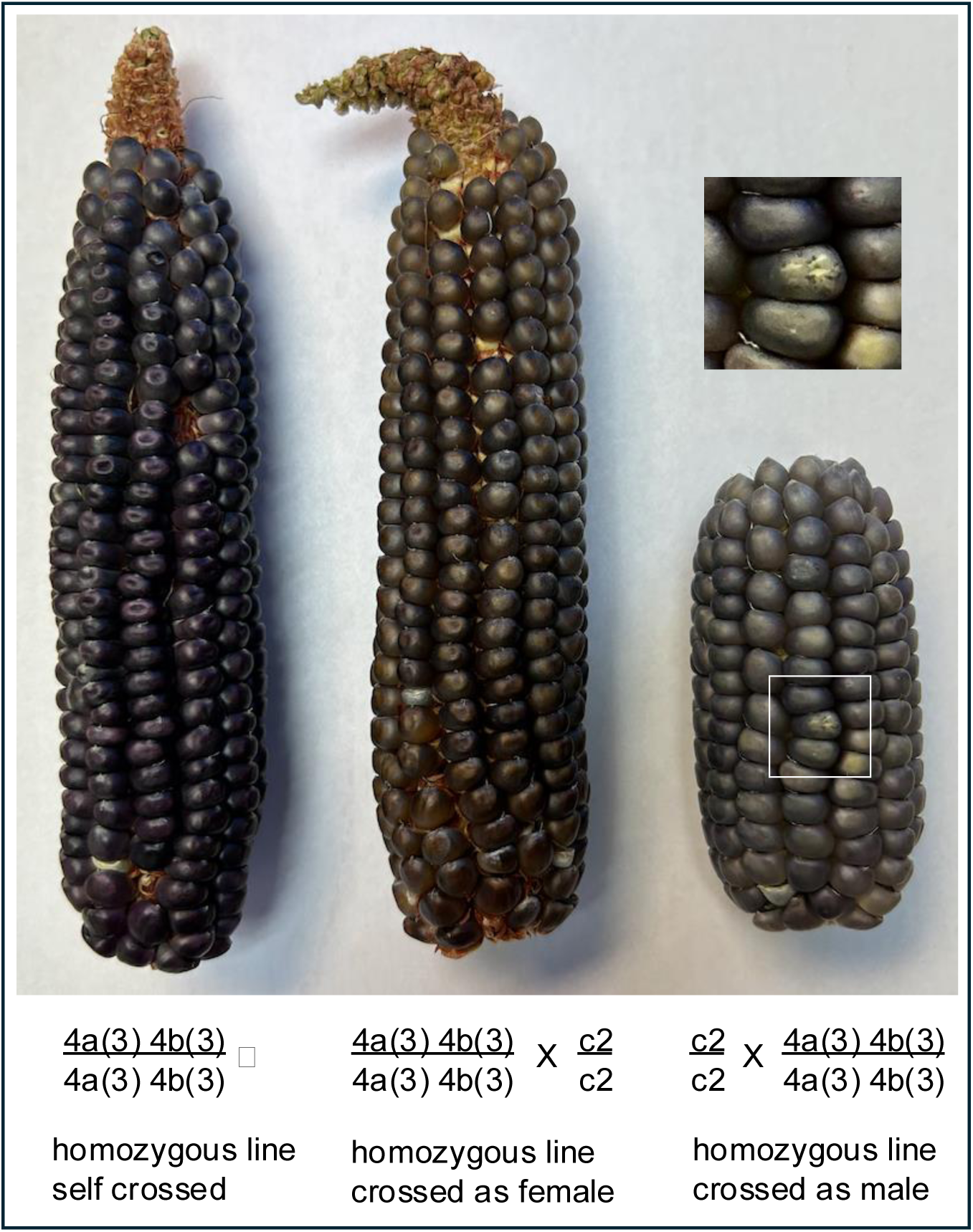
Ears of crosses involving homozygous 4a(3) 4b(3) lines. The inset above the male cross shows a sectored kernel with BFB patterns visible in the endosperm.

**Figure S7:**
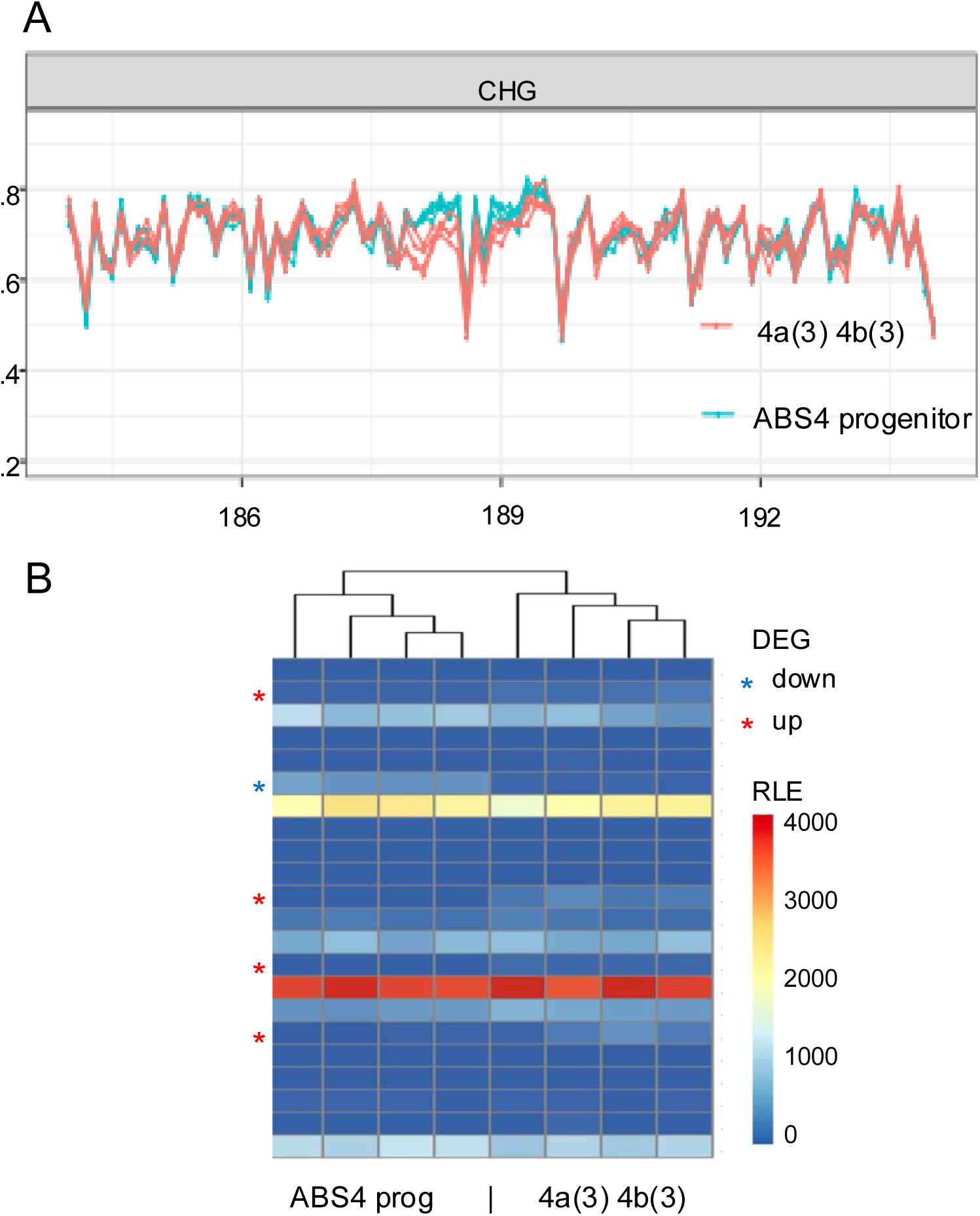
CHG methylation profiles and gene expression analysis of homozygous 4a(3) 4b(3) and ABS4 progenitor lines. **A**) CHG methylation of 4b(3) centromere and its nearby region. Each line indicates one of the four EM-seq samples for each genotype. **B**) Relative Log Expressions (RLEs) of 22 detectable genes in the CENH3-enriched region of 4b(3). Differential expressed genes (DEGs, Wald test) that have significant (Bonferroni adjusted p-value < 0.01) lower expression in 4b(3) are noted with a blue asterisk, while those genes with significant higher expression in the 4b(3) line are noted with a red asterisk. Rows indicate RLEs of one gene. The first four columns are control group ABS4 homozygous plants KD4315.4, KD4315.8, KD4315.2 and KD4315.8. The last four columns are 4a(3) 4b(3) homozygous plants YZ306.6, YZ306.5, YZ306.2 and YZ306.3 (these same eight samples were used for the EM-seq in panel A, with each of the four replicates of each genotype merged).

**Table S1.**
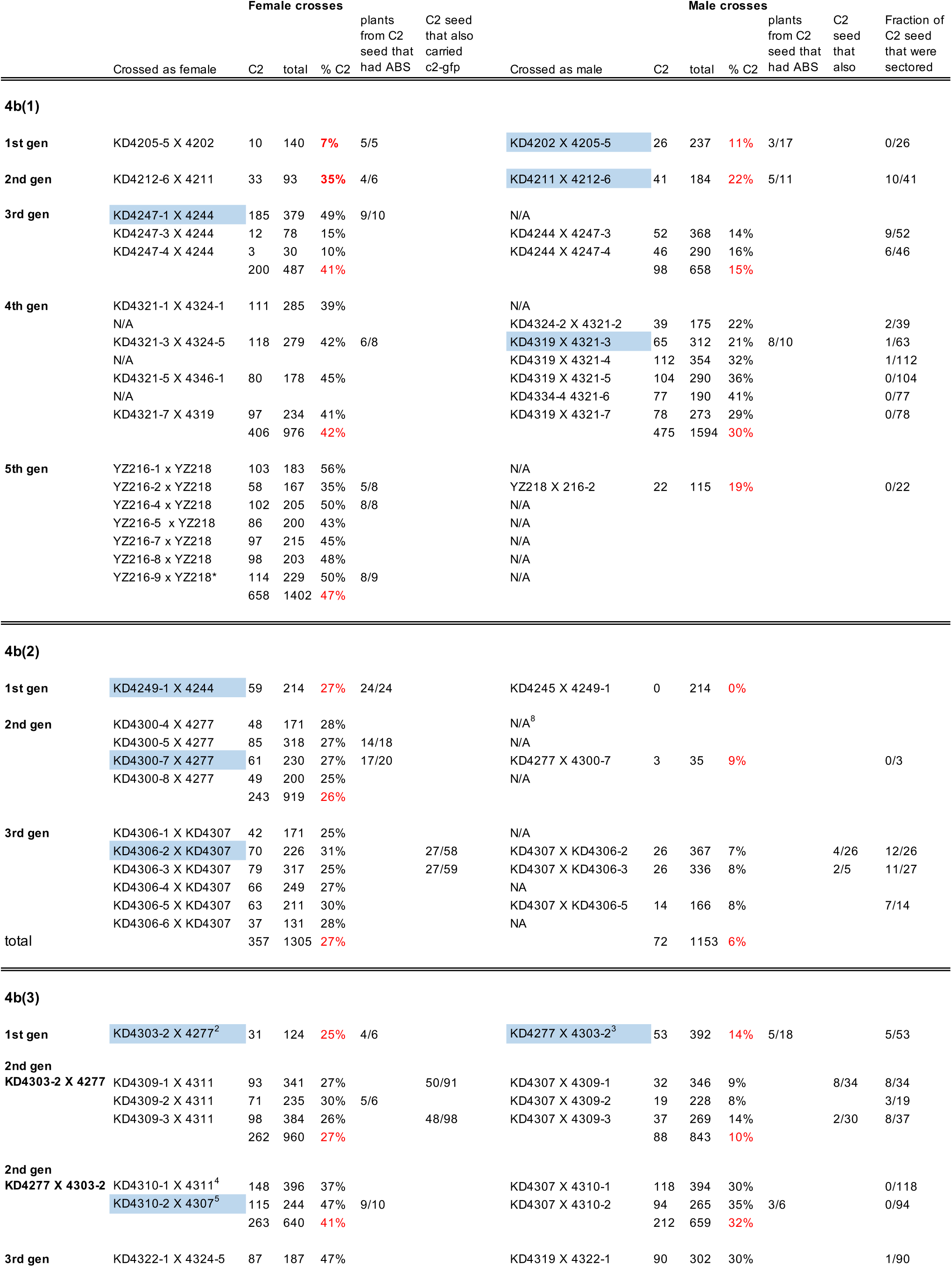
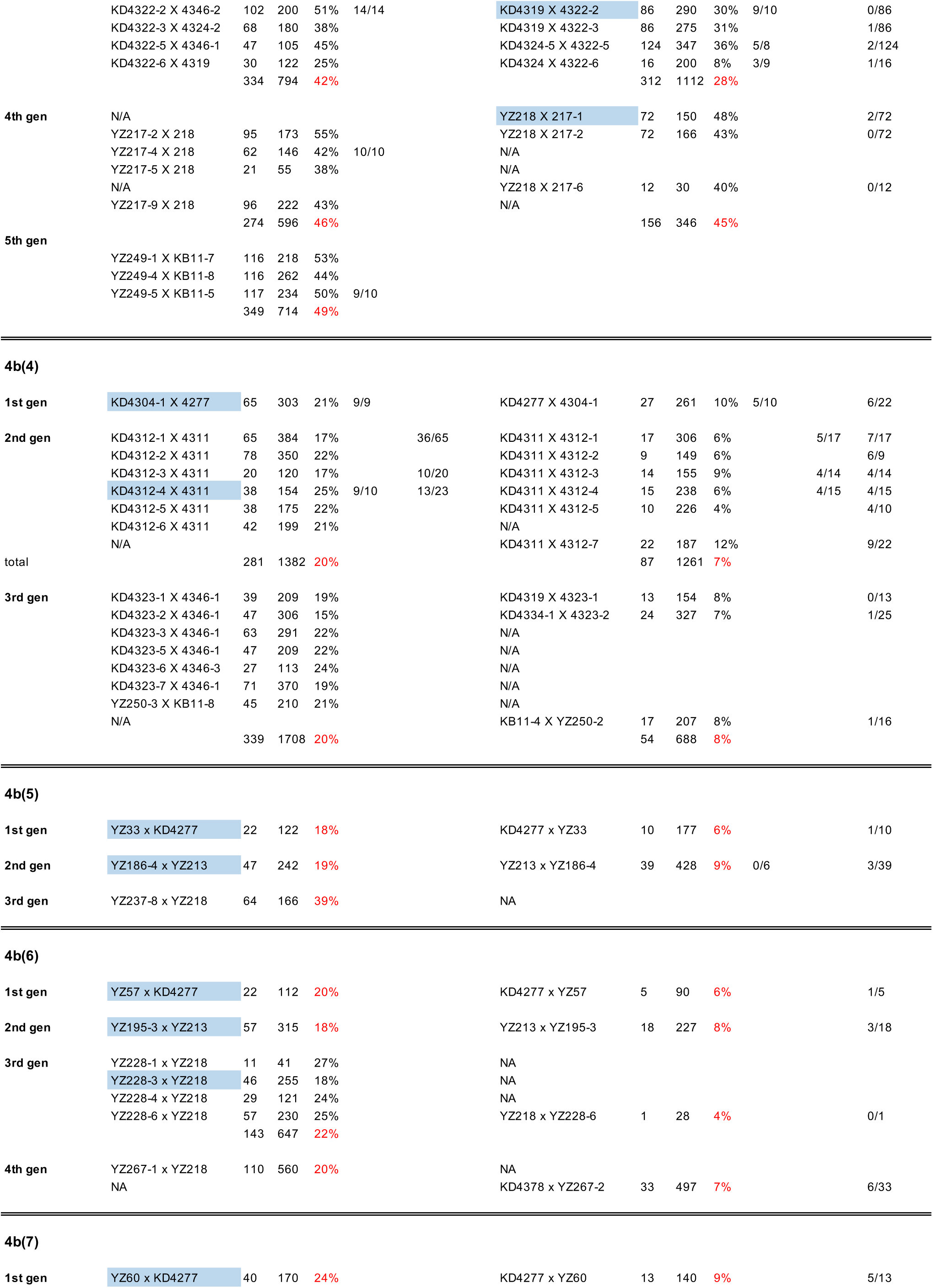
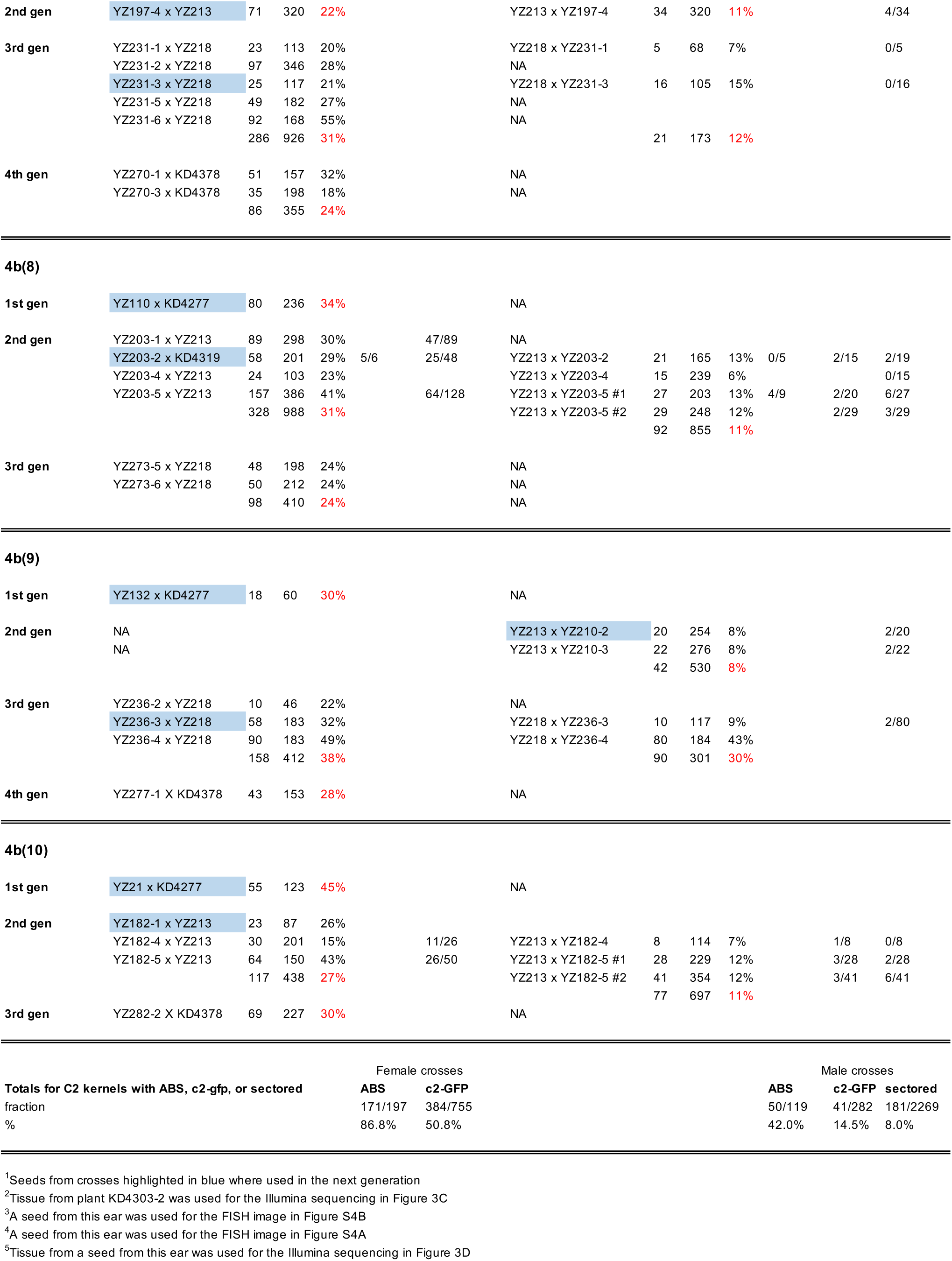
Segregation data for 4b chromosomes^1^.

**Table S2.** Recombination between ABS and C2 in crosses involving partially trisomic 4b(3) lines.

| Female crosses |  |  |  | Male crosses |  |  |  |
| --- | --- | --- | --- | --- | --- | --- | --- |
| KD4309-1 X 4311 | ABS | C2 |  | KD4307 X 4309-1 | ABS | C2 |  |
| Plant 1 | + | + |  | Plant 1 | - | + | recomb |
| Plant 2 | + | + |  | Plant 2 | - | + | recomb |
| Plant 3 | + | + |  | Plant 3 | - | + | recomb |
| Plant 4 | + | + |  | Plant 4 | + | + |  |
| Plant 5 | + | + |  | Plant 5 | - | + | recomb |
| Plant 6 | + | + |  | Plant 6 | - | + | recomb |
| Plant 7 | + | + |  | Plant 7 | + | + |  |
| Plant 8 | + | + |  | Plant 8 | - | + | recomb |
| Plant 9 | + | + |  | Plant 9 | + | + |  |
|  |  |  |  | Plant 10 | + | + |  |
| KD4309-2 x 4307-1 |  |  |  | KD4307 x 4309-2 |  |  |  |
| Plant 1 | + | + |  | Plant 1 | - | + | recomb |
| Plant 2 | + | + |  | Plant 2 | + | + |  |
| Plant 3 | + | + |  | Plant 3 | - | + | recomb |
| Plant 4 | - | + | recomb | Plant 4 | + | + |  |
| Plant 5 | + | + |  | Plant 5 | + | + |  |
| Plant 6 | + | + |  | Plant 6 | - | + | recomb |
| Plant 7 | - | - | loss of both<sup>1</sup> | Plant 7 | - | + | recomb |
| Plant 8 | + | + |  | Plant 8 | - | - | loss of both<sup>1</sup> |
| Plant 9 | + | + |  | Plant 9 | + | + |  |
| Plant 10 | + | + |  | Plant 10 | + | + |  |
| KD4309-3 X 4311 |  |  |  | KD4307 X 4309-3 |  |  |  |
| Plant 1 | + | + |  | Plant 1 | - | + | recomb |
| Plant 2 | + | + |  | Plant 2 | - | + | recomb |
| Plant 3 | - | + | recomb | Plant 3 | + | + |  |
| Plant 4 | + | + |  | Plant 4 | - | + | recomb |
| Plant 5 | - | + | recomb | Plant 5 | + | + |  |
| Plant 6 | - | + | recomb | Plant 6 | - | + | recomb |
| Plant 7 | + | + |  | Plant 7 | - | + | recomb |
| Plant 8 | + | + |  | Plant 8 | + | + |  |
| Plant 9 | + | + |  | Plant 9 | - | + | recomb |
| Plant 10 | + | + |  | Plant 10 | + | + |  |
|  | 24 | 28 |  |  | 13 | 29 |  |
| % recombination |  | 14.3% |  |  |  | 55.2% |  |

<sup>1</sup>Seedlings that lacked both ABS and C2 were scored as non-recombinants.

**Table S3.** Meiosis in homozygous 4a(3)-4b(3) lines.

| Stage | What was scored | Plant number |  |  | totals |
| --- | --- | --- | --- | --- | --- |
|  |  | YZ306-3 | YZ306-10 | YZ306-16 |  |
| Pachytene | Full synapsis of 4a and 4b chromosomes | 31 | 103 | 103 | 237 |
| Diplotene | 4a and 4b paired and visibly separate in cell | 6 | 10 | 12 | 28 |
| Diakinesis | 4a and 4b paired and visibly separate in cell | 1 | 48 | 10 | 59 |
| Metaphase I | Normal metaphase plate | 3 | 5 | 7 | 15 |
| Anaphase I | Normal anaphase segregation | 1 | 14 | 4 | 19 |
| Telophase I | No evidence of micronuclei | 0 | 8 | 0 | 8 |
| Metaphase II | Normal metaphase plate | 0 | 3 | 0 | 3 |
| Anaphase II | Normal anaphase segregation | 3 | 10 | 0 | 13 |
| Telophase II | No evidence of micronuclei | 1 | 20 | 0 | 21 |

**Table S4.** Seed abortion, chromosome loss, and sectoring in crosses of homozygous 4a(3)-4b(3) lines.

| Type of cross | Plant number | Total kernels | Aborted kernels | Kernels w/ purple scutellum | Sectored kernels |
| --- | --- | --- | --- | --- | --- |
| Self and sib crosses |  |  |  |  |  |
| Homozygous 4a(3)-4b(3) self cross | YZ305-1 self | 195 | 4 |  |  |
| Homozygous 4a(3)-4b(3) sib cross | YZ305-2 X sib-1 | 221 | 2 |  |  |
| Homozygous 4a(3)-4b(3) sib cross | YZ305-3 X sib-2 | 128 | 4 |  |  |
| Homozygous 4a(3)-4b(3) self cross | YZ305-25 self | 315 | 2 |  |  |
| Homozygous 4a(3)-4b(3) self cross | YZ305-35 self | 184 | 4 |  |  |
| Homozygous 4a(3)-4b(3) self cross | YZ306-21 self | 274 | 4 |  |  |
| Homozygous 4a(3)-4b(3) self cross | YZ306-26 self | 195 | 1 |  |  |
| total percent |  |  | 1.39% |  |  |
| Female drosses |  |  |  |  |  |
| Homozygous 4(3)-4b(3) X R1-scm2 c2/c2 | YZ305-20 X KD4413 | 93 | 2 | 93 |  |
| Homozygous 4(3)-4b(3) X R1-scm2 c2/c2 | YZ306-27 X KD4413 | 339 | 5 | 339 |  |
| total percent |  |  | 1.62% | 100% |  |
| Male crosses |  |  |  |  |  |
| c2/c2 R1-scm2 X homozygous 4a(3)-4b(3) | KD4413-6 X YZ306-21 | 258 | 3 | 258 | 3 |
| c2/c2 R1-scm2 X homozygous 4a(3)-4b(3) | KD4413-7 X YZ306-21 | 214 | 2 | 214 | 1 |
| c2/c2 R1-scm2 X homozygous 4a(3)-4b(3) | KD4413-8 X YZ306-21 | 203 | 2 | 203 | 1 |
| c2/c2 R1-scm2 X homozygous 4a(3)-4b(3) | KD4413-9 X YZ306-27 | 224 | 4 | 224 | 4 |
| total percent |  |  | 1.22% | 100% | 1.00% |
| Controls |  |  |  |  |  |
| c2/c2 self cross | KD4413-1 self | 276 | 4 |  |  |
| c2/c2 self cross | KD4413-2 self | 246 | 3 |  |  |
| c2/c2 sib cross | KD4413-4 X sib-1 | 220 | 3 |  |  |
| c2/c2 self cross | KD4413-5 self | 245 | 4 |  |  |
| total percent |  |  | 1.42% |  |  |

